# Chromosome-level Subgenome-aware *de novo* Assembly of *Saccharomyces bayanus* Provides Insight into Genome Divergence after Hybridization

**DOI:** 10.1101/2024.03.17.585453

**Authors:** Cory Gardner, Junhao Chen, Christina Hadfield, Zhaolian Lu, David Debruin, Yu Zhan, Maureen J. Donlin, Zhenguo Lin, Tae-Hyuk Ahn

## Abstract

Interspecies hybridization is prevalent in various eukaryotic lineages and plays important roles in phenotypic diversification, adaption, and speciation. To better understand the changes that occurred in the different subgenomes of a hybrid species and how they facilitated adaptation, we completed chromosome-level *de novo* assemblies of all 16 pairs chromosomes for a recently formed hybrid yeast, *Saccharomyces bayanus* strain CBS380 (IFO11022), using Nanopore MinION long-read sequencing. Characterization of *S. bayanus* subgenomes and comparative analysis with the genomes of its parent species, *S. uvarum* and *S. eubayanus,* provide several new insights into understanding genome evolution after a relatively recent hybridization. For instance, multiple recombination events between the two subgenomes have been observed in each chromosome, followed by loss of heterozygosity (LOH) in most chromosomes in nine chromosome pairs. In addition to maintaining nearly all gene content and synteny from its parental genomes, *S. bayanus* has acquired many genes from other yeast species, primarily through the introgression of *S. cerevisiae*, such as those involved in the maltose metabolism. In addition, the patterns of recombination and LOH suggest an allotetraploid origin of *S. bayanus*. The gene acquisition and rapid LOH in the hybrid genome probably facilitated its adaption to maltose brewing environments and mitigated the maladaptive effect of hybridization.

## Introduction

It has generally been believed that hybridization between closely related species often leads to inviability and sterility, a phenomenon known as hybrid incompatibility. The Dobzhansky-Muller (DM) model, which proposes that it results from negative epistatic interactions between genes with different evolutionary histories, is a well-regarded explanation for hybrid incompatibility (Dobzhansky 1982; Price et al. 2010). Hybrid incompatibility can act as a reproductive isolating barrier contributing to speciation (Coyne and Orr 2004). Additionally, reduced fertility in hybrids can result from abnormal chromosome segregation during meiosis if the parental genomes are divergent (Coyne and Orr 2004). Nevertheless, recent studies show that interspecies hybridization is prevalent in major eukaryotic lineages, particularly in angiosperms and yeasts, and it is believed to contribute to adaptation to novel environments (Langdon et al. 2019; Taylor and Larson 2019; Gabaldon 2020; Moran et al. 2021; Suvorov et al. 2022). Given that the exchange of genomic content between species is pervasive, it is important to better characterize the impact of hybridization on evolution of hybrid genomes, which will improve our understanding of the genetic basis underlying the adaptation and divergence of species.

The *Saccharomyces* budding yeast species involved in fermentation of various products is a group of organisms in which hybrids are most commonly found (Langdon et al. 2019; Gabaldon 2020). The allopolyploid genome of *Saccharomyces cerevisiae* has been extensively studied. The ancestral *Saccharomyces* lineage experienced a whole genome duplication (WGD) about 100 million years ago (Wolfe and Shields 1997; Kellis et al. 2004). New evidence suggests that the WGD in the *Saccharomyces* lineage was caused by interspecies hybridization (Marcet-Houben and Gabaldon 2015). Soon after the WGD, there was a period of rapid losses of duplicate genes and only ∼10% of WGD ohnologs survived. The retained WGD duplicates are enriched in genes related to glucose metabolism or rapid growth, such as glycolysis genes (Conant and Wolfe 2007), hexose transporters (Lin and Li 2011), and ribosomal protein genes (Mullis et al. 2019). These studies suggested that the WGD or hybridization event played a significant role in the adaptation of *Saccharomyces* species toward aerobic fermentation (Kellis et al. 2004; Thomson et al. 2005; Conant and Wolfe 2007; Lin and Li 2014) and speciation events (Scannell et al. 2006). These studies improved our understanding of the biological significance of interspecies hybridization in speciation and adaptation.

At the genomic level, questions related to what occurred to the genome after a recent allopolyploidy event, such as the earliest genome rearrangements, the mechanisms of gene loss, recombination between subgenomes, and loss of heterozygosity, are not completely understood (Morales and Dujon 2012). The ancient hybridization events, such as the WGD in the ancestral *Saccharomyces* lineage, may not be useful to address these questions as most duplicate genes have been lost. In addition to the ancient hybridization event, recent interspecific hybridization is prevalent in the *Saccharomyces* lineage as they are used to produce fermented beverages (Langdon et al. 2019). The genomes of these recently generated hybrid genomes may serve as ideal systems to study how genomes evolve after hybridization and contributed to adaptation to specific niches. For instance, *S. pastorianus*, which is an interspecies hybrid between *S. cerevisiae* and *S. eubayanus*, is widely used for brewing lager style beers under low temperature in Europe (Libkind et al. 2011). Some chromosomes in *S. pastorianus* strains may have 5 copies, suggesting its highly aneuploid nature (van den Broek et al. 2015; Gorter de Vries et al. 2017). The chromosome-level assembly for *S. pastorianus* strain CBS 1483, based on MinION long-read sequencing, enables the assembly and exploration of the unstable subtelomeric regions, which contain industrially-relevant genes such as the MAL genes (Salazar et al. 2019).

*Saccharomyces bayanus* is another interspecies hybrid yeast commonly found in industrial brewing environments, but it is viewed as a contaminant in some brewing processes due to the production of undesired byproducts (Rainieri et al. 2003). The taxonomic classification of *S. bayanus* has been a controversial process (Hittinger 2013). Thanks to the discovery of a wild species *S. eubayanus* (Libkind et al. 2011), it is now commonly accepted that *S. bayanus* is a hybrid between *S. uvarum*, and *S. eubayanus* (Perez-Traves et al. 2014; Peris et al. 2014). *S. bayanus* isolates are highly heterogeneous in genetic and metabolic characteristics, probably resulting from many independent hybridization events between *S. eubayanus* and *S. uvarum*, creating many different strains (Rainieri et al. 2006; Libkind et al. 2011; Langdon et al. 2019). Genome sequencing using Illumina has been carried out for over 40 *S. bayanus* strains, such as CBS 380, NCAIM 676, FM1309 and NBRC1948 (Libkind et al. 2011; Almeida et al. 2014; Langdon et al. 2019). Mapping Illumina reads to different *Saccharomyces* species showed that the contributions of genome content from *S. uvarum* and *S. eubayanus* are highly variable among *S. bayanus* strains. Specifically, the genome content deriving from *S. uvarum* ranges from 36.6% to 98.8% (Langdon et al. 2019). In addition, small introgressed regions from *S. cerevisiae* are present in some *S. bayanus* strains (Nguyen et al. 2011). However, due to the limitation of Illumina short reads, these *S. bayanus* genome assemblies are fragmented.

A chromosomal-level subgenome assembly of *S. bayanus* is expected to provide much more detail in the genome evolution following a recent allopolyploidy event. In this study, we sequenced the genome of *S. bayanus* strain CBS 380 (BY20106, IFO11022) using the Nanopore MinION. The strain CBS 380 is the most representative isolate of *S. bayanus*, which has been widely used in many studies (Libkind et al. 2011; Nguyen et al. 2011; Caudy et al. 2013; Perez-Traves et al. 2014). We generated chromosome-level subgenome assemblies based on MinION reads and characterized the evolution of genome structure and gene content. Our results show that *S. uvarum* contributed to over 60% of the hybrid genome. Many chromosomes exhibit mosaic segments of different origins, suggesting multiple recombination events occurred between the two subgenomes after hybridization. Rapid loss of heterozygosity (LOH) following hybridization was also observed, resulting in over 56% of the genome regions becoming homozygous. Introgression from a third species, *S. cerevisiae,* was also detected, contributing to the expansion of maltose metabolism genes in the *S. bayanus* genome. These observations provide detailed examples illustrating how genome evolved immediately after hybridization occurred, improving our understanding of genetic basis of a hybrid species’ survival by overcoming hybrid incompatibility.

## Results

### MinION sequencing, ploidy analysis, and parental inference of *S. bayanus* CBS 380 genome

It is well accepted that *S. bayanus* arose from interspecies hybridization between two closely related *Saccharomyces sensu stricto* yeast species *S. uvarum* and *S. eubayanus* (Figure 1A). To confirm the ploidy levels of the *S. bayanus* CBS 380 strain, we assessed its relative genomic DNA content by fluorescence flow cytometry analysis using a haploid yeast strain *S. bayanus* YJF1450 as a control (Figure 1B). Dual peaks of fluorescence were observed in both strains, with the first peak indicating the DNA content of G1 phase and the second peak showing DNA content after DNA synthesis (G2/M phase). As shown in Fig 1B, the relative genomic DNA content in G1 phase of *S. bayanus* CBS 380 is similar to the G2/M phase the haploid control *S. bayanus* YJF1450, confirming that two sets of chromosomes are present in *S. bayanus* CBS 380.

**Figure 1.**
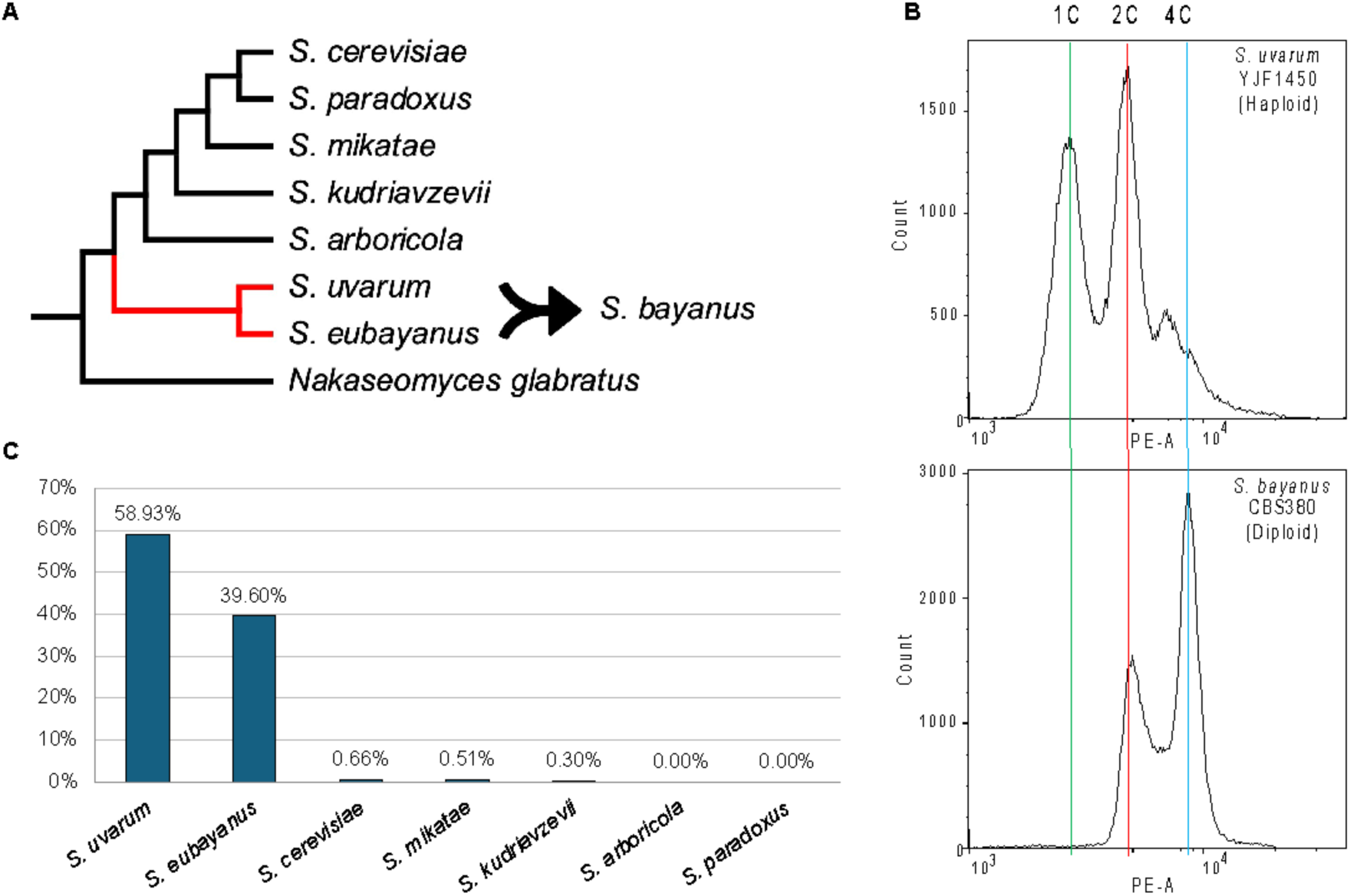
*S. bayanus* CBS 380 has a diploid genome, resulting from a hybridization event between *S. uvarum* and *S. eubayanus*. (A) Schematic illustration of evolutionary relationships among *S. bayanus* and closely related species. (B) Ploidy analysis by flow cytometry of *S. bayanus*. The top histogram shows cell count of *S. uvarum* YJF1450 and the bottom histogram is CBS 380. The x-axis indicates the amount of DNA that is stained by propidium iodide. The green line shows the DNA amount of the G1 phase of the YJF1450 cells (one copy of haploid genome, 1C). The red line shows the G2 phase of the YJF1450 (two copies of haploid genome, 2C) and the G1 phase of the CBS 380 (one copy of diploid genome). The blue line indicates the G2 phase of the CBS 380 (two copies of diploid genomes, 4C). (C) sppIDer results show that most reads from our *S. bayanus* sequencing are mapped to either *S. eubayanus* or *S. uvarum*, confirming the species we sequenced is a hybrid of *S. eubayanus* and *S. uvarum*.

Sequencing of *S. bayanus* CBS 380 with Oxford Nanopore’s MinION yielded 2.2 gigabase pairs (Gb) of data (∼170x coverage), with 2.04 Gb passing quality control (Supplemental Figure S1). Among these, 100 reads exceeded 100 kilobase pairs (Kb) with the longest extending to 158,255 base pairs (bp). We hypothesized that most of our reads would map to the suspected parental species, *S. eubayanus* and *S. uvarum*, while fewer, if any, reads would map to the other more distantly related species. We used sppIDer (Langdon et al. 2018), which maps sequencing reads to the reference genomes of multiple species of interest, to validate the strain sequenced as a hybrid of *S. eubayanus* and *S. uvarum*, and to determine the relative genetic contribution by each parent. The genomes of *S. uvarum* (CBS 7001), *S. eubayanus* (FM 1318), *S. cerevisiae* (BY 4742), *S. mikatae* (IFO 1815), *S. kudriavzevii* (IFO1802), *S. arboricola* (ZP960) and *S. paradoxus* (CBS 432), were used as reference genomes for read mapping. The majority of the reads mapped to *S. eubayanus* and *S. uvarum*, as expected, with 39.6% mapping to *S. eubayanus* and 58.93% mapping to *S. uvarum.* Of the remaining, 5.52% mapped to *S. cerevisiae,* 2.67% to *S. mikatae* and 2.14% to *S. paradoxus* (Figure 1C). This confirms the identity of our sequenced strain as *S. bayanus*, a hybrid of *S. eubayanus* and *S. uvarum*.

### *De novo* assembly and subgenome phasing

Our genome assembly process examined several tools, detailed in the methods section and in Supplemental Table S1, to address the challenges posed by the diploid nature of the target organism. Among the various tools tested, Flye stood out by producing a collapsed-consensus assembly with the highest quality, as reflected in a 96.6% completeness score according to BUSCO analysis. This high score indicates a successful capture of the genomic features we aimed to assemble.

Given the diploid nature of our target organism, we aimed to separately assemble the two subgenomes, diverging from traditional methods that generate a single, collapsed consensus sequence. Using the MinION platform’s long reads, we produced separate and accurate assemblies for each subgenome. Among the methods employed, phasing the Flye collapsed-consensus assembly via the Whatshap pipeline proved the most successful at constructing a high-fidelity diploid genomic representation of *S. bayanus* CBS 380 (Patterson et al. 2015). Post-assembly correction and polishing resulted in a robust genomic structure ready for further analysis (Table 1). To address the inherent complexity of the diploid genome of *S. bayanus* CBS 380, our methodology successfully assembled two distinct subgenomes, technically designated as haplotype-a and haplotype-b, allowing for a sophisticated analysis of the dual genome architecture (Table 1 and Supplemental Table S2). These precise subgenome reconstructions paved the way for gene prediction and other evolutionary insights, with the phased variant calls described in detail in the Materials and Methods.

**Table 1.** Assembly statistics for the *S. bayanus genome* and subgenome grouping. The table details the genomic assembly metrics for the *S. bayanus* species, including the total genome and two subgenomes, Haplotype-a and Haplotype-b, along with the mitochondrial genome. As ancestral subgenomes cannot be directly inferred, homologous chromosomes have been categorized into two hypothetical subgenomes to facilitate analysis.

|  |  | Total genome<br>excluding<br>mtDNA | Subgenome grouping |  | Mitochondrial<br>DNA (mtDNA) |
| --- | --- | --- | --- | --- | --- |
|  |  |  | Haplotype-a | Haplotype-b |  |
| Assembly | Genome Size (bp) | 23,484,151 | 11,829,624 | 11,654,527 | 64,655 |
|  | # of Sequences | 32 | 16 | 16 | 1 |
|  | Largest (bp) | 1,292,201 | 1,292,201 | 1,163,801 | 64,655 |
|  | Smallest (bp) | 208,383 | 217,795 | 208,383 | 64,655 |
|  | Mean (bp) | 733,879 | 739,352 | 728,408 | 64,655 |
|  | N50 (bp) | 912,922 | 912,922 | 919,249 | 64,655 |
|  | GC (%) | 40.1 | 40.1 | 40.1 | 16.23 |

|  | N Count | 300 | 100 | 200 | 0 |
| --- | --- | --- | --- | --- | --- |
| Annotation | Genes | 11,545 | 5,737 | 5,808 | 20 |
|  | CDS | 12,789 | 6,318 | 6,471 | 8 |

### Genome annotation

Evidence based prediction and annotation of protein-coding genes for each subgenome/haplotype of *S. bayanus* CBS 380 was carried out using the GALBA pipeline (Bruna et al. 2023). The pipeline is perfectly suited to our use case, given its capability to leverage high-quality protein sequences from closely related species. The output revealed a total of 11,547 protein-coding genes identified across both haplotypes, with 5,737 genes in haplotype a and 5,808 genes in haplotype b (Table 1 and Figure 2). The variation in gene count between the two haplotypes is in direct proportion to their chromosomal lengths.

**Figure 2.**
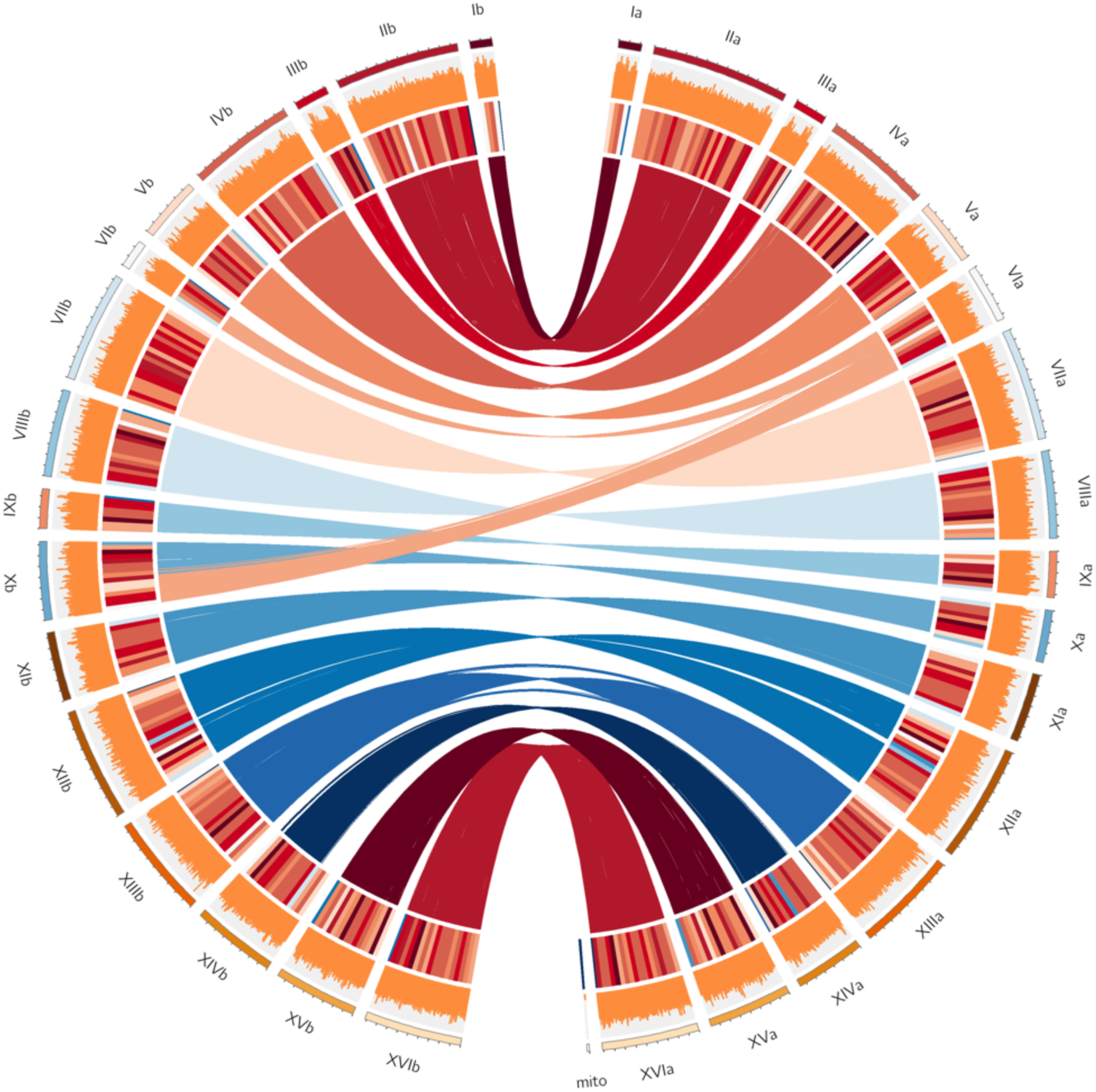
Circos plot representing the 16 pairs of chromosomes of the *S. bayanus* genome, showing different genomic features across four concentric circles. The outermost circle represents the karyotype of the *S. bayanus* genome for two haplotypes, with the right part representing haplotype-a and the left part representing haplotype-b. The second outermost circle represents the GC content on each chromosome of the genome. The third circle provides information on gene density within the chromosomes. The innermost circle highlights synteny blocks between haplotypes, illustrating regions of genetic similarity and divergence between haplotype-a and haplotype-b.

The completeness of the genome annotation was accessed by BUSCO based on saccharomycetes_odb10 database, which indicates a high degree of completeness (2095 of 2137, 98%). As the BUSCO analysis was based on both haplotype assemblies, most genes are expected to have two copies. As a result, 86.2% of genes (1,843) were classified as duplicates, while only 11.8% (252) were identified as unique single-copy genes. Only 13 genes (0.6%) from the saccharomycetes_odb10 gene set were absent from our predicted list. The genome annotation, CDS, and protein sequences are available at https://github.com/BioHPC/Saccharomyces-bayanus.

Functional annotation of the predicted genes was conducted using Eggnog-mapper (Cantalapiedra et al. 2021), which assigned key functional information, such as descriptions of biological functions, orthologous genes in *S. cerevisiae*, Gene Ontology, KEGG pathway and Pfam domains, to 10,985 genes, accounting for 95.1% of the total identified genes (Supplemental Table S3). The combination of functional annotation and BUSCO assessments confirms that our annotation results are comprehensive, providing a solid foundation for our further analysis.

### Inference of parental genomic regions

To identify the major genomic events that have occurred in the *S. bayanus* genome since hybridization, including recombination, chromosomal rearrangements, and loss of heterozygosity, we first used two approaches to determine the origin of genomic regions in the hybrid genome. Our first approach is based on BLAST searches of non-overlapping blocks of 5,000 bp for every chromosome against the genomes of *S. eubayanus* and *S. uvarum* (see Methods and Materials). In brief, the origin of each genomic block was determined by its best hit of BLAST search. As illustrated in Figure 3A, each haplotype chromosome contains regions that originated from both *S. uvarum* and *S. eubayanus*, suggesting the recombination between the two orthologous chromosomes of the two subgenomes, creating mosaic chromosomes composed of genomic regions of heterozygous origins. However, the proportions of each subgenome vary substantially across different chromosomes. For instance, segments of *S. eubayanus* origin make up 81% of Chr IVa, whereas they make up only 14% of Chr IVb. In addition, nine of the 16 chromosome pairs have a high degree of homozygosity, meaning that the genomic origin and recombinants are very similar between haplotypes a and b, showing that heterozygosity was quickly lost after hybridization.

**Figure 3.**
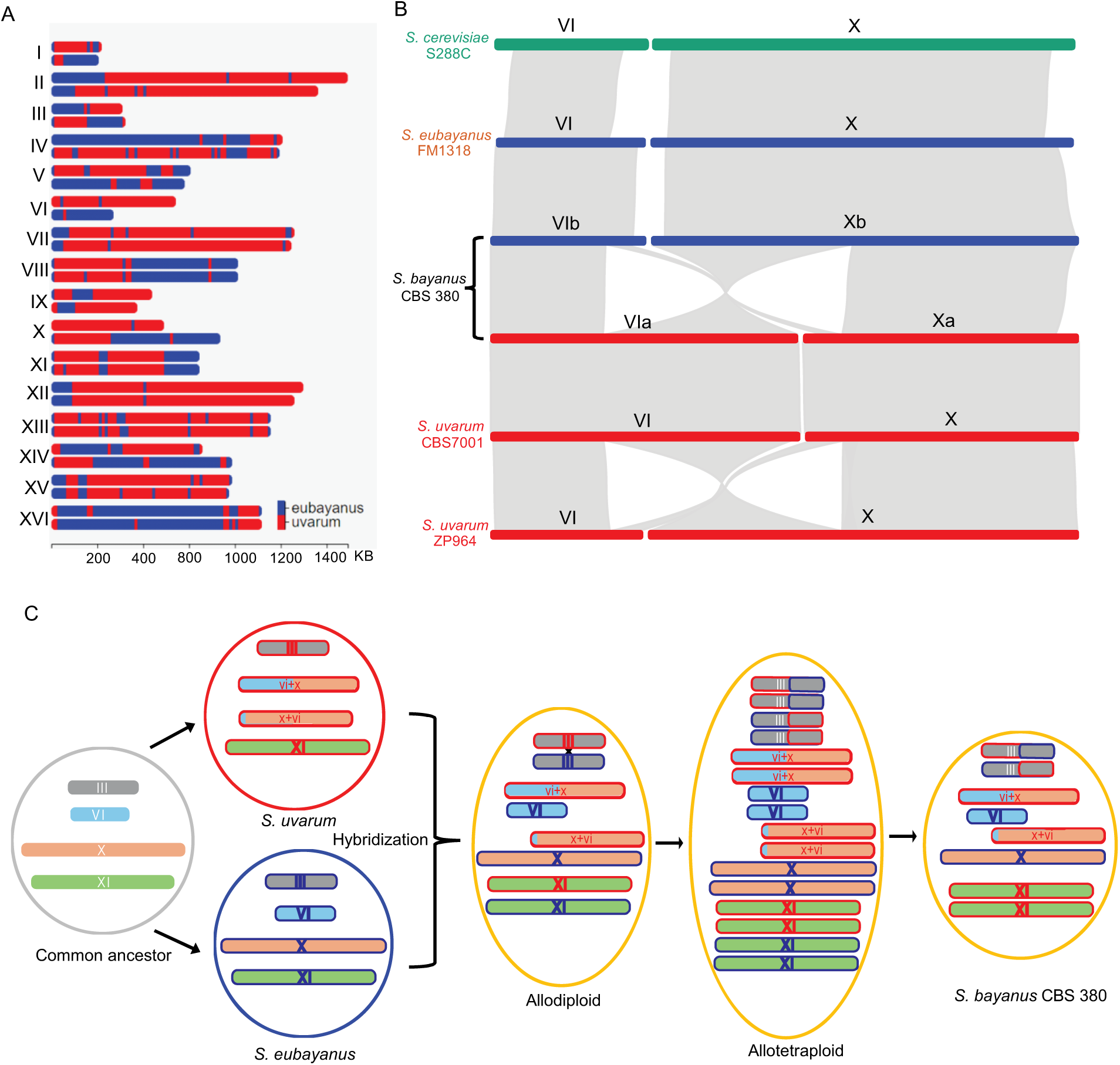
The origin and evolution of *S. bayanus* chromosomes. (A) Origin of genomic regions of each chromosome in the *S. bayanus* genome based on BLAST searches of non-overlapping 5,000 bp blocks. Genomic Regions that originated from *S. eubayanus* are shown in blues, while regions inherited from *S. uvarum* are shown in red. (B) Synteny block of Chromosome VI and X between *S. cerevisiae, S.ubayanus,* both *S. bayanus* haplotypes, *S. uvarum* strain CBS 7001, *S. uvarum* strain ZP964. (C) An evolutionary model of S*. bayanus* chromosomes. For simplification purposes, only four chromosomes are shown, representing different patterns of chromosome inheritances. Translocation between Chr VI and X occurred in *S. eubayanus* prior to its hybridization with *S. bayanus*. Recombination and whole genome duplication occurred in the hybrid *S. bayanus* genome. Subsequent genome reduction by chromosome losses, created some heterozygous chromosomes, such as Chr III, and some homozygous chromosomes, such as Chr XI.

To confirm the robustness of the results based on the BLAST method, we employed an approach based on the degree of divergence of synonymous sites in protein-coding regions, as it is generally assumed that synonymous mutations are selectively neutral (see Methods and Materials). To summarize, the method first compared rates of synonymous substitution (*Ks*) between the two alleles of *S. bayanus* and then between orthologous genes from the three species. We then determined if an allele in *S. bayanus* is more similar to its orthologous gene in *S. uvarum* or in *S. bayanus*. We identified a total number of 5,497 orthologous groups from the three species using OrthoFinder (Emms and Kelly 2019) (Supplemental Materials). The distribution of *Ks* between all pairs of alleles in *S. bayanus* shows two distinct peaks (Supplemental Figure S2). The left peak, which consists of lower *Ks* values, represents sequence divergence between two homozygous alleles (alleles originated from one single parental genome). In contrast, the right peak contains higher *Ks* values that were obtained from two alleles originated from different parental genomes (heterozygous origins). Consistently, the distribution of *Ks* values between orthologous genes between the two parental genomes largely overlaps with the right peak of *Ks* values in *S. bayanus* (Supplemental Figure S2). Next, we calculated *Ks* values between an allele from *S. bayanus* and its orthologous gene in *S. uvarum* (*Ksu*) and *S. eubayanus* (*Kse*) respectively. The origin of each allele was then determined by comparing *Ksu* and *Kse* to the two *Ks* peaks. The results of genomic origin obtained based on our *Ks* method are highly consistent with the BLAST method (Supplemental Figure 3), supporting the occurrence of multiple recombination events on most chromosomes and rapid loss of heterozygosity after interspecies hybridization.

### A model of allotetraploid origin of *S. bayanu*s

We found distinct differences in the lengths of chromosomes VI and X between the two subgenomes (haplotypes) of *S. bayanus* CBS380 (Figures 2 and 3A, Supplemental Table S2). Specifically, Chr VIa is ∼277k bp longer than Chr VIb (544k vs. 267k), while Chr Xa is ∼244k bp shorter than Chr Xb. Our analysis of syntenic regions between the two haplotypes shows that a significant portion of Chr VIa has syntenic regions to Chr Xb. These observations suggest that a translocation occurred between Chr VI and X. Next, we sought to determine whether the translocation was from Chr VI to Chr X or visa versa, and whether the translocations occurred in the parent genomes or after hybridization. The answers to these questions are key to better understanding how hybrid *S. bayanus* arose, and the mechanism by which heterozygosity is rapidly lost on most chromosomes.

To investigate the direction and timing of the translocation event, we conducted a detailed examination of all available genomes of *S. uvarum* and *S. eubayanus* strains from the NCBI database. Our results showed that the patterns of chromosome lengths in all *S. eubayanus* strains were consistent with those observed in *S. cerevisiae*, i.e., a short chromosome VI and a long chromosome X (Figure 3B). In contrast, heterogenous lengths of Chr VI and Chr X are observed among S. *uvarum* strains (Figure 3B). For instance, S. *uvarum* ZP964 has similar lengths of Chr VI and Chr X to those of *S. eubayanus* and *S. cerevisiae*. In contrast, *S. uvarum* CBS 7001 has a much longer Chr VI, but much shorter Chr X, similar to the haplotype a in *S. bayanus* (Figure 3B). Gene collinearity analysis of the two chromosomes among these species further showed that the translocation occurred only once in the lineage of *S. uvarum* CBS 7001 prior to hybridization*. S. bayanus* inherited translocated Chr VI and Chr X from a parental species that is closely related to *S. uvarum* CBS 7001. These results also support that the translocation was generated by exchanging ∼270 KB segment at the left end of Chr X with ∼30 KB region at the right end of Chr VI (Figure 3B).

Our analysis of origins of genomic regions in *S. bayanus* demonstrates that only seven pairs of chromosomes maintained heterozygous status, and loss of heterozygosity occurred to other chromosome pairs (Figure 3A). Several genetic mechanisms have been proposed to explain the LOH after hybridization, such as whole-genome duplication followed by chromosome loss, duplication or loss of individual chromosomes, and gene conversion (Marcet-Houben and Gabaldon 2015; Wolfe 2015; Wertheimer et al. 2016). Duplication and loss of individual chromosomes often result in chromosomal aneuploidies. However, we did not observe obvious chromosomal aneuploidies in *S. bayanus* based on read depth of most, if not all, chromosomes. In addition, the track length of gene conversion is usually limited, which is not supported by our observations that the track length of LOH covers almost entire chromosomes. Furthermore, the locations of recombination events are very similar between haplotypes in most chromosomes, such as Chr VIII, Chr XI, and Chr XII (Figure 3A). Based on these observations, it is mostly parsimonious to propose that the hybrid alloploid genome may have undergone duplication without cell division (non-disjunction), resulting in a temporary allotetraploid genome. Subsequence loss of chromosomes may have occurred to the allotetraploid genome, resulting in haploid status (Figure 3C). Therefore, the allotetraploid origin of *S. bayanus* is similar to the evolutionary history of *S. pastorianus* (Dunn and Sherlock 2008; Nakao et al. 2009; Libkind et al. 2011).

### Evolution of gene content after hybridization and expansion of genes involved in maltose metabolism

To better understand the evolution of gene content after hybridization, we carried out further analyses on the 5,497 orthologue groups (OG) in *S. bayanus* and its parental species *S. uvarum* and *S. eubayanus*. A total number of 5,412 unique OGs are present in the two parent genomes. 5,389 of them (99.6%) are also present in *S. bayanus*, suggesting that gene loss in *S. bayanus* is very limited after hybridization (Figure 4A). Interestingly, 85 OGs (195 genes) are only present in the genome of *S. bayanus*. Given that ∼5% of MinION reads were specifically mapped to *S. cerevisiae*, we speculated that these genes were likely originated from introgression events of *S. cerevisiae* as proposed in a previous study (Nguyen et al. 2011).

**Figure 4.**
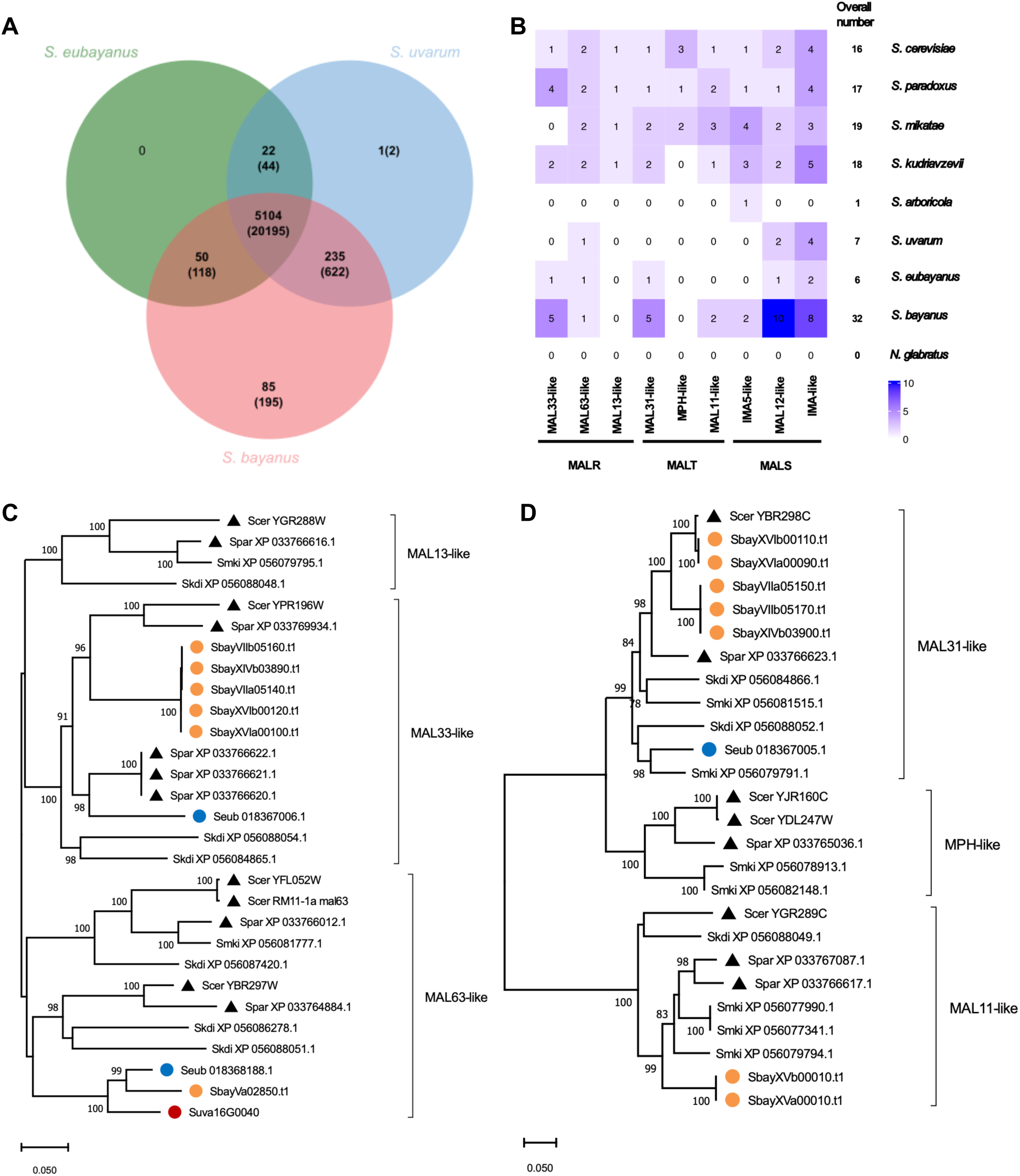
The evolution of gene content in the hybrid genome of *S. bayanus* CBS 380. (A) A Venn diagram showing the numbers of shared and species-specific orthologous groups (OGs). (B) Evolutionary changes of the three MAL gene families in nine OGs in the Saccharomyces sensu stricto members and *Nakaseomyces glabratus*. Increased gene copy numbers were observed in each MAL gene family in *S. bayanus*. (C) A phylogenetic tree of the MALR gene family suggests introgression of MALR genes in *S. bayanus*. (D) A phylogenetic tree of the MALT gene family suggests that all MALT genes in *S. bayanus* were likely acquired by introgression events from other species.

Among the 5,104 OGs present in all three species, 4,065 OGs (79.6%) exhibit a 1:1:2 ratio, indicating conservation of gene copy numbers in all three species. It is worth noting that 200 of these OGs contain more than two copies of genes in the diploid genome of *S. bayanus*, suggesting expansion of gene copy number in *S. bayanus*. Nine of these expanded OGs are involved in maltose/maltotriose utilization (Figure 4B). *S. bayanus* is mainly found in brewing environments and maltose is the most abundant sugar (∼60%) in brewer’s wort (Magalhaes et al. 2016). Therefore, the expansion of genes involved in maltose or maltotriose utilization (MAL genes) might have facilitated the adaptation of *S. bayanus* to maltose-rich environments.

MAL genes were classified into three families based on their functions, including maltose transporter (MALT), enzymes that break down maltose (MALS), and genes that regulate the expression of the pathway (MALR). These genes are often organized into clusters and located near the ends of chromosomes (subtelomeric). To elucidate the origins and expansion of MAL genes in *S. bayanus*, we first identified all MAL genes in *S. bayanus* and eight other *Saccharomyces* species. The total numbers of MAL genes vary substantially among non-hybrid species, ranging from 0 in *Nakaseomyces glabratus*, to 19 in *S. mikatae* (Figure 4B). The diploid genome of *S. bayanus* contains a significantly higher number of MAL genes (32 in total) compared to 7 in *S. uvarum* and 6 in *S. eubayanus* (Figure 4B), suggesting a significant expansion in copy numbers of MAL genes in *S. bayanus* after hybridization. This is particularly noticeable that the MALS gene family was increased from 3 and 6 in their parental species genomes to 20 in *S. bayanus*.

To infer the origin and mechanism of MAL gene expansion in *S. bayanus*, we carried out phylogenetic analyses for each of the three MAL families using their amino acid sequences (Figure 4C-D, Supplemental Figure S4). Based on the tree topology, we found that the majority of the MAL genes in *S. bayanus* were not inherited directly from its parental genomes. Instead, these genes seem to be likely acquired through introgression from other species, mostly from *S. cerevisiae*, followed by multiple gene duplication events. For example, 6 copies of MALR genes are present in *S. bayanus*. Only one MALR gene (MAL63-like) is group with *S. eubayanus*, while the other five form a well-supported clade that is closely related to MAL33-like genes in *S. cerevisiae* and *S. paradoxus* (Figure 4C), suggesting multiple rounds of gene duplication to MAL33-like genes after acquisition of MAL33-like genes from ancestral *S. cerevisiae* or *S. paradoxus*. Similarly, none of the seven MALT genes was grouped with either *S. eubayanus* or *S. uvarum* (Figure 4D). In the case of MALS genes, a total number of 20 MALS genes are found in *S. bayanus*, and only 8 of them appear to be originated from *S. eubayanus*, and expanded by gene duplication. Similar to other *Saccharomyces* species, most MAL genes in *S. eubayanus* also form clusters and reside in subteleomeric regions (Supplemental Figure S5).

We noticed that none of *S. bayanus* MAL genes were inherited from *S. uvarum* (Figure 4C-D, Supplemental Figure S4). It suggests that there was a preferred retention of MAL genes inherited from *S. eubayanus* or a preferred loss of MAL genes inherited from *S. uvarum*. Given that *S. uvarum* has contributed over 60% of the genetic makeup of *S. bayanus*, the strong exclusion of *S. uvarum* MALs genes in the *S. bayanus* genome were unlikely due to random events. One possibility is that MAL genes from *S. uvarum* might imposed selective disadvantages under maltose-rich brewing environments. Future studies on the growth effects of *S. uvarum* MAL genes may provide new insights into the biased retention of MAL genes.

## Discussion

We present the first chromosome level subgenome assembly and annotations of the hybrid yeast, *S. bayanus* (CBS 380) which will serve as an excellent reference for future studies of this important yeast and other yeast strains. The assembly was completed using only Oxford Nanopore technology on a single MinION flow cell. Thus, we show the utility of high read depth sequencing, that is available for moderate costs using this technology. We assessed the assemblies from fifteen different *de novo* assembly pipelines, all run on relatively modestly equipped computer workstations, and concluded that the Flye method outperformed the others in producing an assembly with the fewest contigs and high N50 scores. The successful application of the GALBA pipeline allowed for high-fidelity annotation of the two subgenomes, revealing a total of 11,547 protein-coding genes and confirming the completeness of our genome assembly with a high BUSCO score. This type of sequencing can be carried out in most laboratories without previous sequencing experience or high-performance computational resources.

Through a dual approach involving BLAST searches and synonymous site divergence, we traced recombination events and chromosomal rearrangements that describe the history of *S. bayanus* after hybridization. In particular, the discovery of mosaic chromosomes with heterozygous origins in *S. uvarum* and *S. eubayanus* speaks to the dynamic evolutionary past of this species. Our data suggest a rapid loss of heterozygosity, which can be attributed to multiple genetic mechanisms, including a proposed transient polyploid phase and selective chromosome loss. This model not only parallels the evolutionary history of related species such as *S. pastorianus*, but also provides a plausible explanation for the observed genome organization.

Although the majority of genes in *S. bayanus* maintained their copy numbers, large copy number variations were found in the three MAL families. In addition to direct transmission from parent species, our phylogenetic analysis suggested that many of them were acquired from other yeast species, mostly from *S. cerevisiae*. Subsequent gene duplication events on MAL genes further increased its copy number. It is reasonable to postulate that the introgression and duplication of MAL genes provide selective advantages in maltose-rich brewing environments.

Despite *S. bayanus* inheriting most chromosomal segments from *S. uvarum*, none of the MAL genes in *S. bayanus* were traced to its *S. uvarum* progenitor. It is unlikely due to random loss of *S. uvarum* copies. One possibility is that *S. eubayanus* MALs genes are more efficient or provide more selective advantages than those of *S. uvarum*. Further studies can be performed to examine the functional differences of these MAL genes in maltose metabolism between the two species, which could provide new valuable information for improving industrial brewing using maltose-rich materials.

## Materials and Methods

### Yeast strain, growth condition, genomic DNA isolation

*Saccharomyces bayanus* CBS 380’s cells were grown on YPD medium (1% yeast extract, 2% peptone, and 2% glucose) at 30 °C for 16 hours. Extraction of high molecular weight genomic DNA (HMW gDNA) from *S. bayanus* cells was carried out by following a protocol described by Denis et al (Denis E 2018). In brief, *S. bayanus* cell wall was first lysed with Zymolyase (MP Biomedicals). Spheroplasts were then collected and resuspended in SDS buffer with RNase A. Proteins were precipitated and removed with potassium acetate and centrifugation. The supernatants were used to precipitate DNA with isopropanol. DNA pellet was then washed with 70% ethanol and dissolved in TE buffer. The quality and quantity of the extracted DNA were determined using Qubit (Invitrogen). HMW gDNA was sheared into 20kb fragments using g-TUBE (Covaris Inc).

### Determination of ploidy

We performed a flow cytometry analysis to determine the ploidy of the *S. bayanus* CBS 380 following the protocol (Todd et al. 2018). We also used a haploid *S. uvarum* strain YJF1450 (MATα hoΔ::NatMX, derived from CBS 7001, a gift from J. Fay lab at Rochester University) as a control. Briefly, yeast cells were grown to log-phase (OD = 0.3) in YPD medium on a shaker platform at 30 °C by rotation at 225 RPM. Then, cells were fixed in 70% ethanol at 4 °C overnight and then sonicated to separate cells. After RNase A (0.5 mg/ml) treatment for 2 hours, the cells were stained with 25 μg/ml of propidium iodide at 4 °C overnight. Finally, the stained cells were analyzed using BD Accuri C6 Plus and the data were analyzed in FlowJo v10.8.1.

### MinION library preparation and sequencing

HMW gDNA were then used to prepare MinION sequencing library using the Nanopore Rapid Sequencing Kit (SQK-RAD004) following the manufacturer’s instruction. Briefly, the sample mix was prepared with 7.5 µl template DNA (∼2µg) and 2.5 µl fragmentation mix and incubated at 30°C for 1 min and then at 80 °C for 1min. 1 µl Rapid Adapter was added to the sample mix and incubated for 5 min at room temperature. Priming mix was prepared by adding 30 μl of Flush Tether and Flush Buffer. The priming mix was loaded into the flow cell via the priming port. Sequencing mix was prepared with DNA sample mix and was loaded to the flow cell via the SpotON sample port.

### Adapter removal

Porechop v0.2.4 (Wick et al. 2017) was used for adapter identification and removal using default thresholds. In all, 179,725 reads had adapters trimmed from their start (15,472,707 bases removed), and 778 reads were split based on middle adapters. (Supplemental Figure 1). A full list of commands and parameters is available in the Supplemental Materials.

### Genome assembly, post-assembly correction, and genome polishing

Draft collapsed-consensus assemblies were generated using Canu v2.2 (Koren et al. 2017), Flye v2.9 (Kolmogorov et al. 2019), Wtdbg2 v2.5 (Ruan and Li 2020), NECAT v0.0.1 (Chen et al. 2021), SMARTdenovo v1.0.0 (Liu et al. 2021), NextDenovo v2.5.0 (NextOmics 2021), Raven v1.8.0 (Vaser et al. 2017), and Ra v0.2.1 (Vaser 2019), with both uncorrected and Canu corrected and trimmed reads (Supplemental Table 1). These methods were executed on a general workstation-level computer (36 cores and 128GB memory), demonstrating the feasibility of ONT-based de novo assembly for small genomes in modestly equipped laboratories.

### Complete subgenome-aware de novo genome assembly

Given the diploid nature of our target organism, we aimed to generate a diploid-level representation of each chromosome. We employed long-read sequencing to facilitate the generation of full-length, phased haplotype *de novo* assemblies, using a suite of assembly tools, as detailed below and in the Supplemental Material.

#### Haplotype-aware de novo genome assembly

We experimented with haplotype-aware assembly methods such as Flye (with haplotype preservation enabled) (Kolmogorov et al. 2019), Shasta (Shafin et al. 2020), Phasebook (Luo et al. 2021), and CanuTrio (which organizes reads into haplotype-specific bins before assembly) (Koren et al. 2017). These approaches did not yield high-quality assemblies that were both contiguous and reflective of the expected genome size, leading to their exclusion from analysis.

#### Phasing-based diploid genome assembly

To tackle the complexities of *S. bayanus* CBS 380’s diploid genome, we undertook a phasing-based assembly strategy, leveraging the long reads generated from Oxford Nanopore’s MinION platform. Prior to phasing, the purge_dups pipeline was used to remove haplotype duplication in the primary assemblies (Guan et al. 2020). To construct a phased diploid genome assembly, we first called variants using Claire (Zheng et al. 2022). The variant calls were processed through the WhatsHap pipeline which exploits the connectivity between heterozygous variants within individual reads to generate phased haplotypes (Patterson et al. 2015). To generate a haplotype-specific genomic representation, we used BCFtools ‘consensus’ followed by WhatsHap manual. This allowed us to extract the separate FASTA representations for each haplotype, effectively translating the phased information into a coherent, usable format for further analysis. BUSCO was used to assess the assembly’s completion (Simao et al. 2015). A comprehensive list of commands and parameters, along with the phased variant calls, are accessible in the Supplemental Materials, offering a resource for future genetic and evolutionary studies.

#### Genome correction and polishing

For assembly correction and polishing, the raw ONT sequencing reads were split via the ‘whatshap split’ subcommand to segregate the set of unmapped reads according to their haplotypes. This generated two distinct FASTQ files, each corresponding to one of the haplotypes identified within the sample. The assembled contigs were then passed to a series of correction and polishing steps to enhance their accuracy, utilizing Racon (v1.4.3) (Vaser et al. 2017) and Medaka (v1.9.1) (https://github.com/nanoporetech/medaka) for error correction and sequence improvement. This correction process was executed separately for each haplotype, utilizing their respective reads. A total of four iterative rounds of correction were iteratively performed with Racon for each haplotype. This cycle involved mapping the haplotype-resolved reads to the assembled contigs using Minimap2 (using the ONT-specific ‘-x map-ont’ option), followed by Racon-based correction to refine assembly quality progressively. After completing the Racon correction cycles, a final round of polishing was conducted using Medaka. This step uses a neural network-based approach to correct consensus sequence errors, further enhancing the accuracy of the assembled haplotypes.

### Genome annotation

We employed the GALBA pipeline to annotate protein-coding genes for the assembled nuclear genome (Bruna et al. 2023). Specifically, we used amino acid sequences from *Saccharomyces cerevisiae*, *Saccharomyces uvarum*, and *Saccharomyces eubayanus* as inputs. These protein sequences were aligned to both subgenomes of *S. bayanus* using the Miniprot (Li 2023), followed by gene annotation using AUGUSTUS (Stanke et al. 2006). The output GTF files were processed using AGAT (https://github.com/NBISweden/AGAT) for format cleaning and conversion. The completeness of the gene annotation was evaluated using the BUSCO (version: 5.5.0) (Simao et al. 2015), employing the saccharomycetes_odb10 database for assessment. For functional annotation of predicted genes, we utilized the web version of eggnog-mapper (Cantalapiedra et al. 2021) to upload the *S. bayanus* protein files. All other parameters were retained as default settings.

### Ancestral inference of *S. bayanus* using a BLAST-based approach

To infer the ancestral parentage of the hybrid yeast, we conducted a comparative genomic analysis using a custom script that performs local BLAST (Camacho et al. 2009) homology searches (see Supplemental Materials). The hybrid yeast genome was segmented into consecutive, non-overlapping chunks of 5,000 base pairs, which were then individually compared against the genomes of the two parental strains using the BLAST algorithm. This approach allowed for the identification of the closest matching regions between the hybrid and each parent genome, based on the highest bitscore values obtained from the BLAST results. The bitscore, serving as a measure of sequence similarity, was selected as the primary criterion for parental inference. A score threshold of 100 bits was set to distinguish between significant and non-significant matches, thereby facilitating the identification of the most probable ancestral parent for each genomic segment of the hybrid yeast.

#### Inference of gene origin in *S. bayanus* based on similarity of synonymous sites

To delineate the evolutionary lineage of genes within *S. uvarum*, *S. eubayanu*s, and *S. bayanus*, we initially conducted an OrthoFinder (Emms and Kelly 2019) analysis on the coding sequence (CDS) datasets of these species to identify orthologous genes. Subsequently, alignments were performed utilizing PRANK (Loytynoja 2021) with codon model. Following alignment, we employed KaKs_Calculator (Zhang 2022) (version 3.0) to compute the synonymous substitution rate (*Ks*). The derived Ks values were then utilized to categorize gene lineage based on their similarity, providing insights into the genetic heritage of the genes of *S. bayanus*.

### Comparative Genomic Analysis and Evolutionary Study of the MAL Gene family in *S. bayanus*

To elucidate the evolutionary relationships among *Saccharomyces bayanus* and closely related species, we analyzed coding sequence (CDS) datasets for *S. cerevisiae*, *S. paradoxus*, *S. mikatae*, *S. kudriavzevii*, *S. arboricola*, *S. uvarum*, *S. eubayanus*, and *Nakaseomyces glabratus*. Using OrthoFinder (Emms and Kelly 2019), we identified orthogroups to enable a comparative genomics study. Adopting the protocols from (Brown et al. 2010) and (Baker et al. 2015), we identified genes belonging to the maltose utilization (MAL) gene families. Sequence alignment was conducted with MAFFT using the L-INS-i strategy (Katoh and Standley 2013). Phylogenetic trees were generated using FastTree v2.1 (Price et al. 2010). The evolutionary history was inferred using the Neighbor-Joining method (Saitou and Nei 1987). The original tree is shown. The percentage of replicate trees in which the associated taxa clustered together in the bootstrap test (1000 replicates) are shown next to the branches (Felsenstein 1985). The tree is drawn to scale, with branch lengths in the same units as those of the evolutionary distances used to infer the phylogenetic tree. The evolutionary distances were computed using the Maximum Composite Likelihood method and are in the units of the number of base substitutions per site. These trees were visualized and refined with MEGA11 (Tamura et al. 2021).

## Acknowledgments

C.G., C.H. D.D. and T.A were supported by the National Science Foundation (NSF) under Grant No. 1564894 and Z. L was supported by NSF under Grant No. 1951332. M.J.D receives funding from the National Institute of Allergy and Infectious Diseases of the National Institutes of Health under award number R01AI123407. We thank Dr. Justin Fay for providing *S. uvarum* YJF1450 strain for ploidy analysis.

## Data Availability

Sequencing and genome assembly data generated in this work have been deposited at the NCBI repository under the BioProject accession PRJNA741321. Annotations and supplemental materials are available at https://github.com/BioHPC/Saccharomyces-bayanus.

## Author Contributions

T.A. and Z.Lin conceived the idea. T.A, Z.Lin, and M.J.D. supervised this study. Z.Lu isolated DNA, prepared libraries and performed Nanopore sequencing. Y.Z. performed flow cytometry. C.G., J.C, C.H., and D.D. analyzed the data. All authors wrote the manuscript and approved the final version of the manuscript.

## Supplemental Materials

**Supplemental Figure S1:**
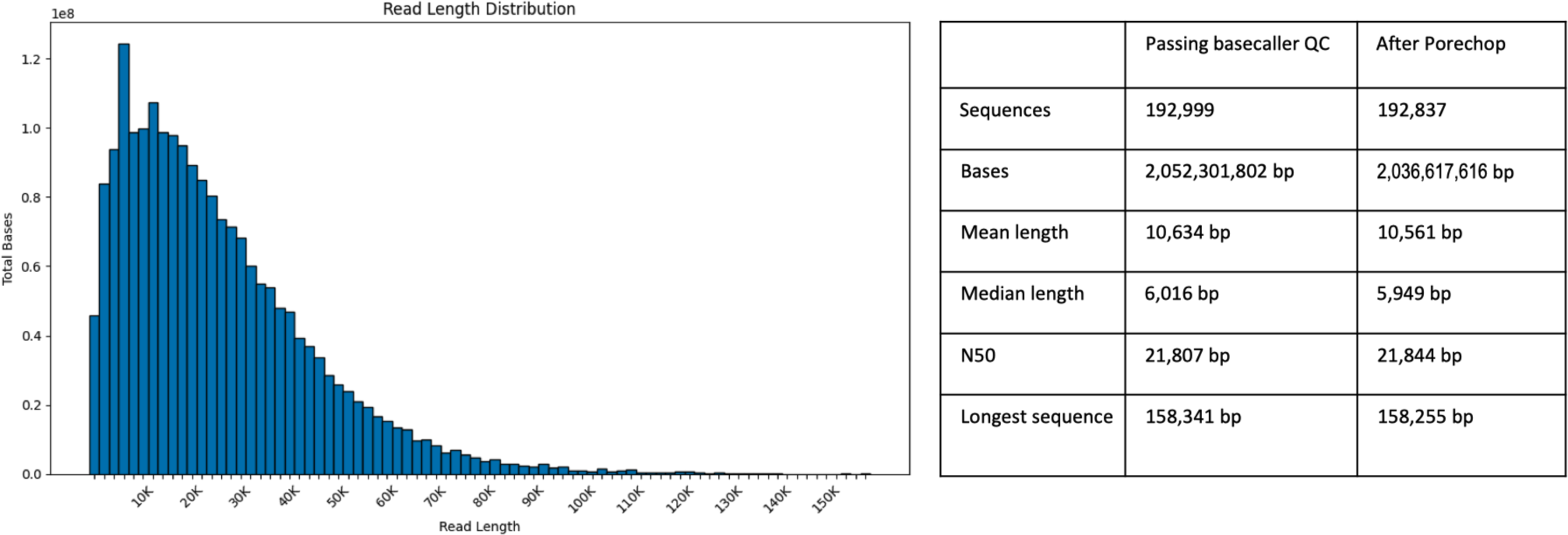
Overview of *S. bayanus* CBS 380 genome sequencing results using Oxford Nanopore’s MinION. The sequencing effort generated 2.2 Gb of data, achieving approximately 170-fold coverage. After quality control, 2.04 Gb of data were retained. Notably, the sequencing run produced 100 reads surpassing 100 Kb, with the longest read measuring 158,255 bp.

**Supplemental Table S1:** Comparative analysis of genome assembly tools used for *S. bayanus* CBS 380. This table summarizes the performance of eight different assembly algorithms, including Flye, NextDeNovo, WTDBG, Ra, NECAT, SmartDeNovo, Canu, and their optimized versions (denoted by an asterisk), with an emphasis on addressing diploidy complexities.

| Assembler | Flye | NextDeNovo | WTDBG | Ra | NECAT | SmartDeNovo | Canu | SmartDeNovo* | Flye* |
| --- | --- | --- | --- | --- | --- | --- | --- | --- | --- |
| Length (bp) | 14,635,841 | 11,870,454 | 11,811,864 | 12,017,687 | 15,428,960 | 11,578,645 | 16,115,554 | 12,391,933 | 13,538,319 |
| Total seq. # | 38 | 17 | 21 | 24 | 32 | 16 | 49 | 21 | 62 |
| Max seq. | 1,286,589 | 1,293,072 | 2,164,800 | 1,101,538 | 1,293,344 | 1,293,262 | 1,292,830 | 1,101,177 | 1,285,351 |
| Min seq. | 2,346 | 122,189 | 17,265 | 8,314 | 59,794 | 61,853 | 10,281 | 59,343 | 2,074 |
| Med seq len. | 301,095 | 773,390 | 462,238 | 528,214 | 442,886 | 780,948 | 213,365 | 610,360 | 84,583 |
| N50 (length) | 647,014 | 816,895 | 813,059 | 785,083 | 646,646 | 919,172 | 612,488 | 787,374 | 785,325 |
| N50 (ctg #) | 9 | 6 | 5 | 7 | 9 | 6 | 9 | 7 | 7 |
| BUSCO | C:96.6%[S:74.5%<br>D:22.1%],F:2.3%,M:1.1% | C:96.6%[S:74.5%<br>D:22.1%],F:2.3%,M:1.1% | C:96.6%[S:74.5%<br>D:22.1%],F:2.3%,M:1.1% | C:96.6%[S:74.5%<br>D:22.1%],F:2.3%,M:1.1% | C:96.6%[S:74.5%<br>D:22.1%],F:2.3%,M:1.1% | C:96.6%[S:74.5%<br>D:22.1%],F:2.3%,M:1.1% | C:96.6%[S:74.5%<br>D:22.1%],F:2.3%,M:1.1% | C:96.6%[S:74.5%<br>D:22.1%],F:2.3%,M:1.1% | C:96.6%[S:74.5%<br>D:22.1%],F:2.3%,M:1.1% |

\* Ran with Canu-corrected and trimmed reads

**Supplemental Table S2:** Our methodology successfully assembled two distinct subgenomes, technically designated as haplotype-a and haplotype-b, allowing for a sophisticated analysis of the dual genome architecture. This table shows the different size of each chromosome in subgenomes.

| Chr. | Haplotype-a | Haplotype-b |
| --- | --- | --- |
| I | 217,795 | 208,383 |
| II | 1,292,201 | 1,163,801 |
| III | 306,526 | 319,184 |
| IV | 1,007,622 | 1,000,830 |
| IX | 433,831 | 370,629 |
| V | 601,693 | 581,353 |
| VI | 544,124 | 266,821 |
| VII | 1,055,917 | 1,042,311 |
| VIII | 817,384 | 817,171 |
| X | 488,404 | 732,591 |
| XI | 646,725 | 646,334 |
| XII | 1,096,630 | 1,057,074 |
| XIII | 959,152 | 958,029 |
| XIV | 664,155 | 791,919 |
| XV | 784,543 | 778,848 |
| XVI | 912,922 | 919,249 |
| Mt | 64,655 |  |

**Supplemental Data D1**: A list of f 5,497 orthologous groups and gene members from the three species are available at https://github.com/BioHPC/Saccharomyces-bayanus.

**Supplemental Figure S2:**
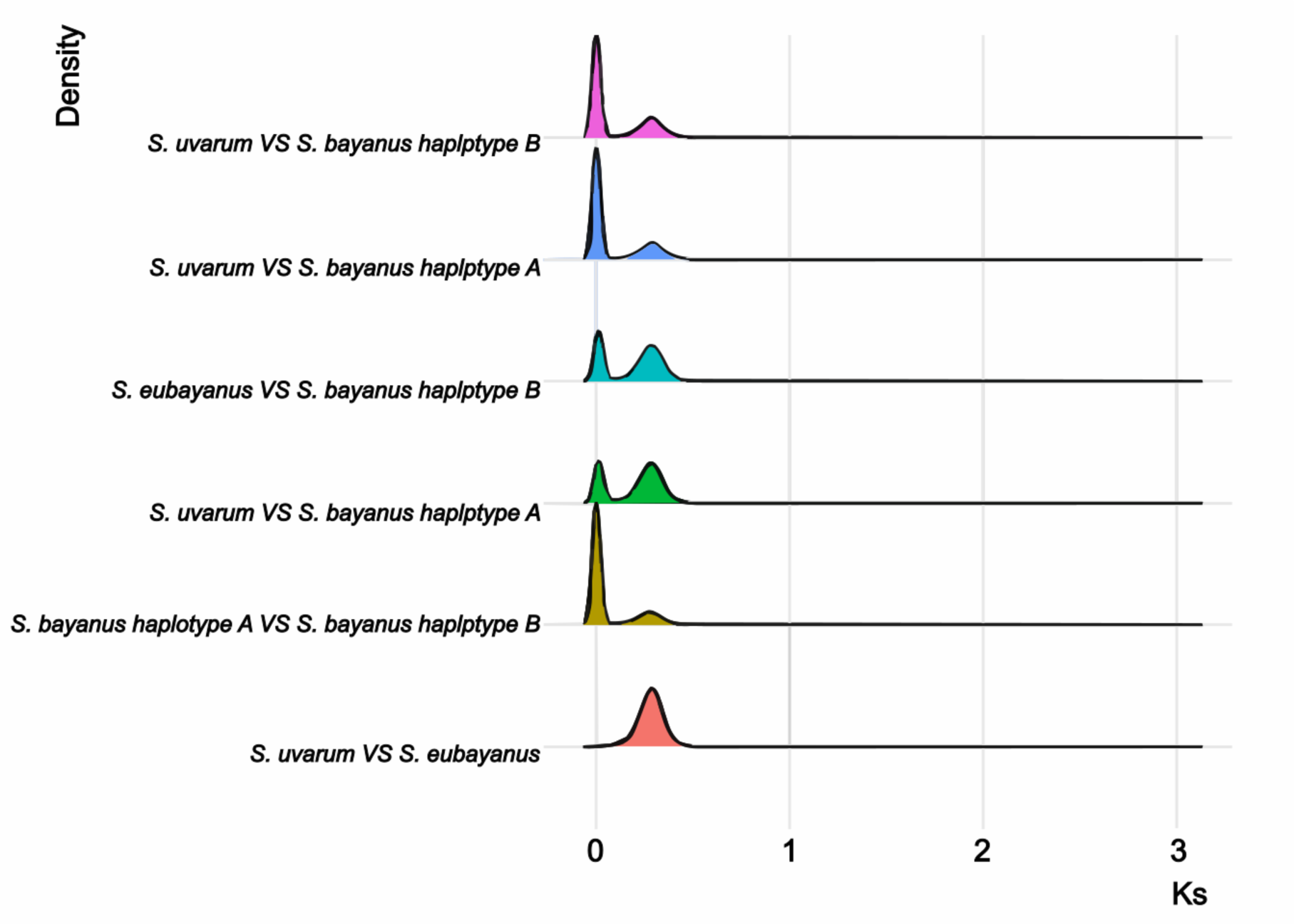
Distribution of *Ks* values.

**Supplemental Figure S3:**
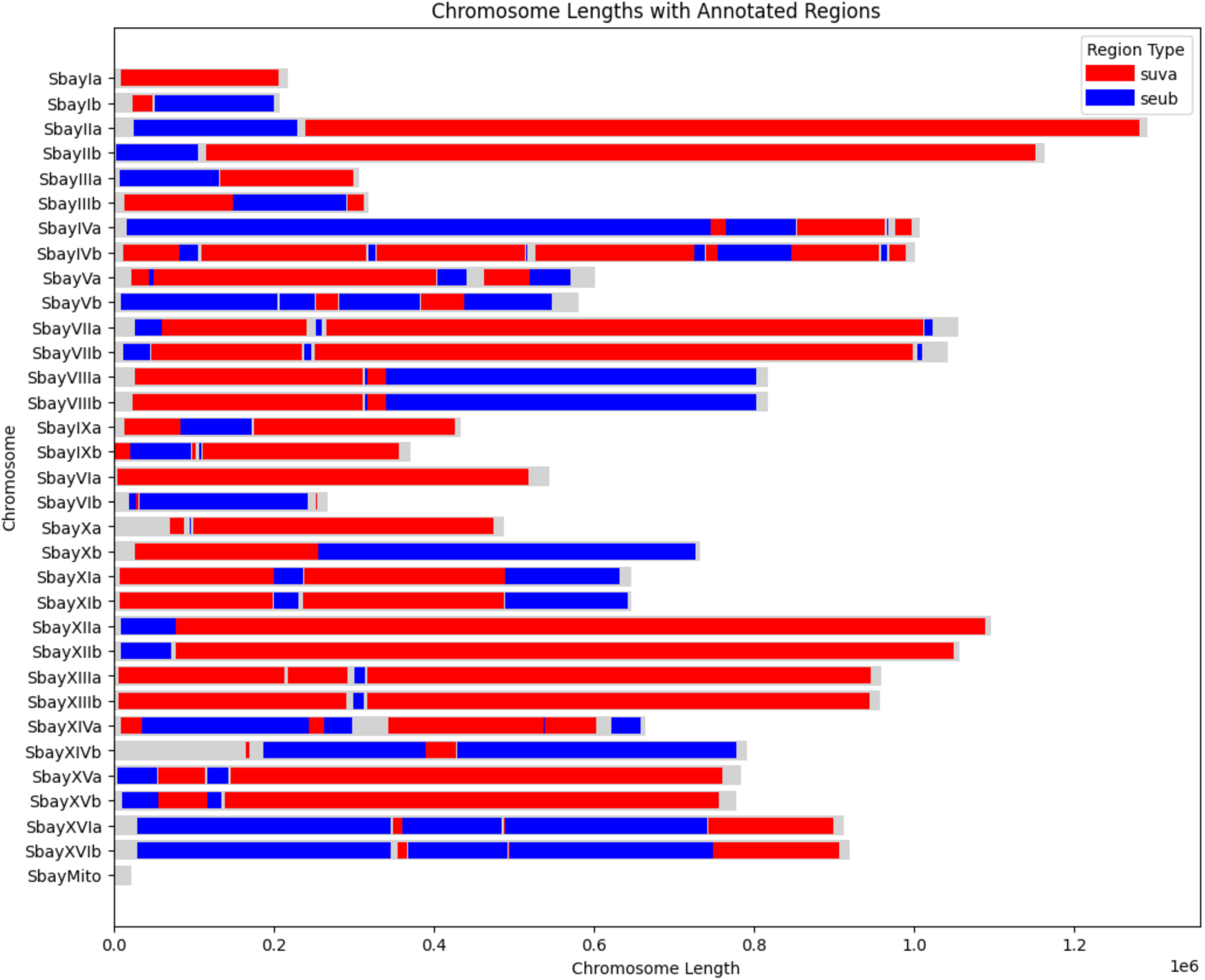
Genomic origin based on gene *Ks* values.

**Supplemental Figure S4:**
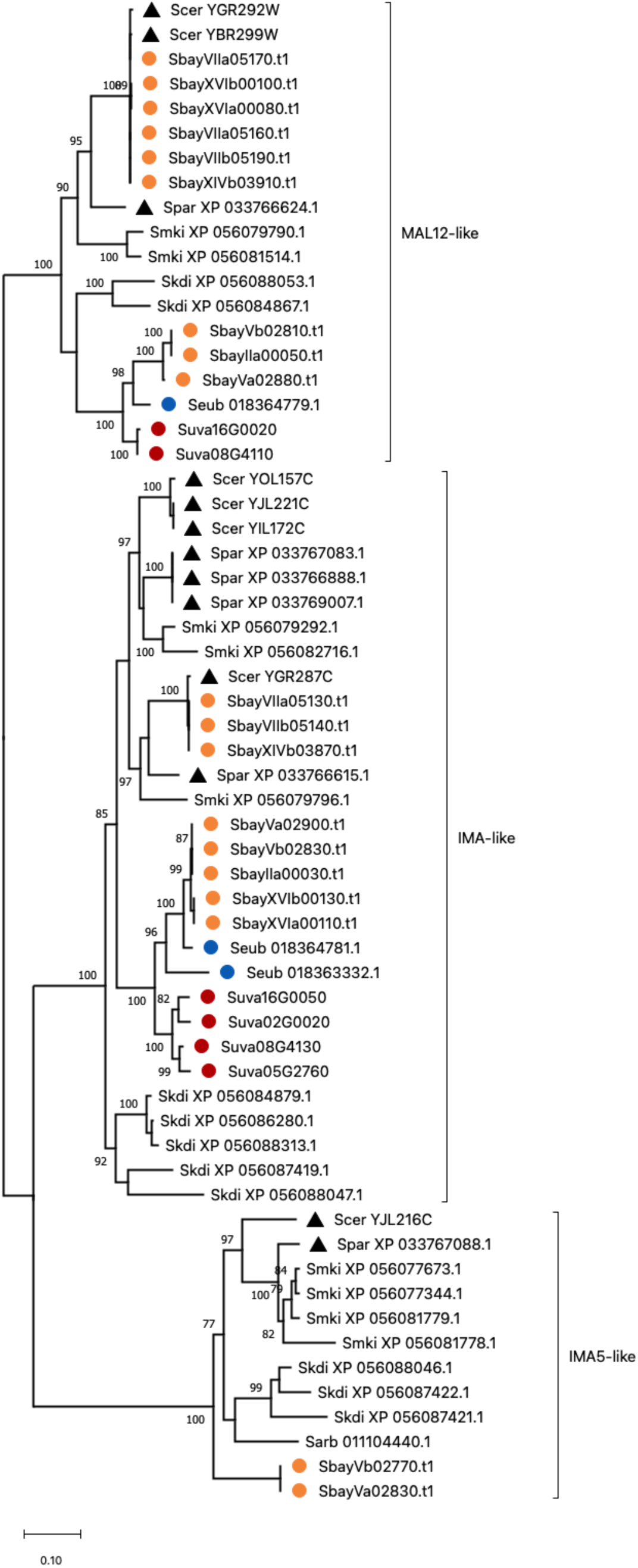
A phylogenetic tree of the MALS gene family.

**Supplemental Figure S5:**
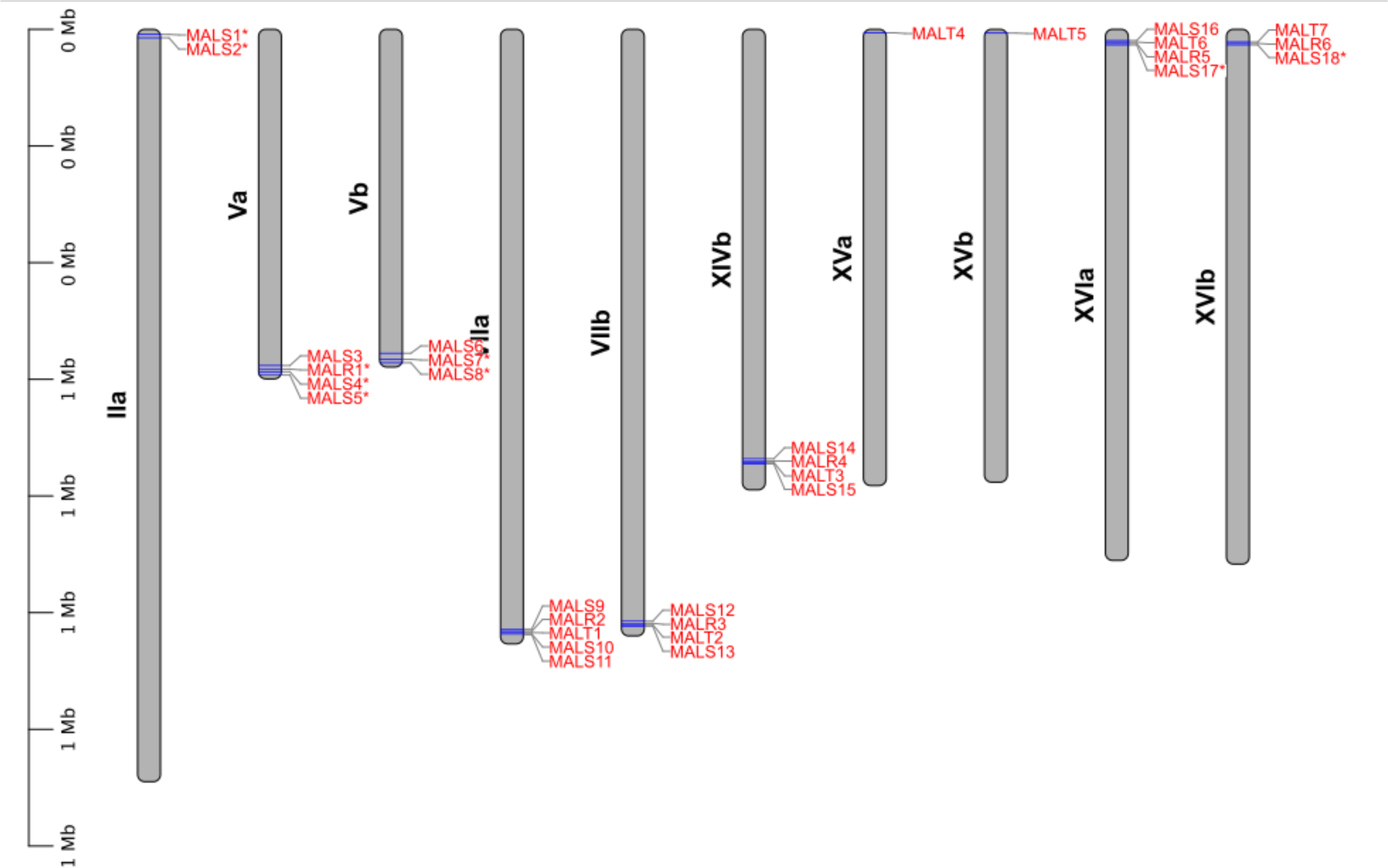
The location of MAL gene families. Gene with star means inherited from *S. eubayanus*.

